# Factors that affect the rates of adaptive and non-adaptive evolution at the gene level in humans and chimpanzees

**DOI:** 10.1101/2021.05.05.442740

**Authors:** Vivak Soni, Adam Eyre-Walker

## Abstract

The rate of amino acid substitution has been shown to be correlated to a number of factors including the rate of recombination, the age of the gene, the length of the protein, mean expression level and gene function. However, the extent to which these correlations are due to adaptive and non-adaptive evolution has not been studied in detail, at least not in hominids. We find that the rate of adaptive evolution is significantly positively correlated to the rate of recombination, protein length and gene expression level, and negatively correlated to gene age. The correlations remain significant when each factor is controlled for in turn, except when controlling for expression in an analysis of protein length; and they also remain significant, or marginally significant, when biased gene conversion is controlled for. However, the positive correlations could be an artefact of population size contraction. We also find that the rate of non-adaptive evolution is negatively correlated to each factor, and all these correlations survive controlling for each other and biased gene conversion. Finally, we examine the effect of gene function on rates of adaptive and non-adaptive evolution; we confirm that virus interacting proteins (VIPs) have higher rates of adaptive and lower rates of non-adaptive evolution, but we also demonstrate that there is significant variation in the rate of adaptive and non-adaptive evolution between GO categories when removing VIPs. We estimate that the VIP/non-VIP axis explains about 5-8x more of the variance in evolutionary rate than GO categories.

## Introduction

There is substantial variation in the rate of evolution between different genes within a genome; some genes, such as those coding for histones, evolve very slowly, whereas many genes involved in immunity evolve rapidly (Clark et al. 2003; Chimpanzee Sequencing and Analysis Consortium, 2005; Nielsen et al. 2005; Sackton et al. 2007; Obbard et al. 2009). The reasons for this variation have been extensively studied and a number of factors appear to influence or be correlated to the rate of protein evolution including function (e.g. Proschel et al. 2006; Haerty et al. 2007; Obbard et al. 2009), mutation rate (Taddei et al. 1997; Tenaillon et al. 1999; Giraud et al. 2001; Denamur & Matic, 2006; Lynch et al. 2016), recombination rate (RR) (Hill & Robertson 1966; Marais & Charlesworth, 2003), gene expression (Pal et al. 2001; Subramanian & Kumar, 2004; Wright et al. 2004; Lemos et al. 2005) and protein length (Zhang, 2000; Lipman et al. 2002; Liao et al. 2006). Correlations with other factors, such as essentiality, appear to be less clear (Hurst & Smith, 1999). Any one of these patterns could be due to adaptive or non-adaptive evolution, but the relative roles of these two different evolutionary processes have rarely been studied.

At the functional level, genes involved in immunity, tumor suppression, apoptosis and spermatogenesis have been shown to have higher rates of adaptive evolution in hominids (Clark et al. 2003; Nielsen et al. 2005; Chimpanzee Sequencing and Analysis Consortium, 2005). Particularly striking is the amount of adaptive evolution that appears to occur in virus-interacting genes, which appear to account for 30% of all adaptive substitutions in hominids, whilst these genes only constitute 13% of the proteome by length (Enard et al. 2016). In *Drosophila* it has been shown that male-biased genes, such as testes specific genes, have higher rates of adaptive evolution (Proschel et al. 2006; Haerty et al. 2007), as do genes involved in immunity (Sackton et al. 2007; Obbard et al. 2009). The dominant role of VIPs in hominid adaptive evolution begs the question of whether there is variation between other categories of genes, and how much of the variation in the rate of adaptive evolution is partitioned between the VIP and non-VIP categories. The role of gene function in determining non-adaptive evolution has not been addressed in detail.

The rate of protein sequence evolution has been shown to be correlated to gene expression, with highly expressed genes having lower rates of protein evolution in both eukaryotes (Pal et al. 2001; Subramanian & Kumar, 2004; Wright et al. 2004; Lemos et al. 2005) and prokaryotes (Rocha & Danchin, 2004). Moutinho et al. (2019) has shown that this correlation is due to both adaptive and non-adaptive evolution in *Drosophila* suggesting that gene expression constrains the rate of adaptive substitution as well as the effect of purifying selection. In *Arabidopsis* the correlation with expression seems to be largely associated with non-adaptive evolution (Moutinho et al. 2019). The role of gene length has also been studied, with several studies showing that smaller genes evolve more rapidly (Zhang, 2000; Lipman et al. 2002; Liao et al. 2006). Again, this appears to be due to both adaptive and non-adaptive evolution, in *Drosophila* species, but possibly only due to non-adaptive evolution in *Arabidopsis* (Moutinho et al. 2019).

Genes differ not only in function, expression, and length, but also in age (Lynch, 2002; Daubin & Ochman, 2004; Tautz & Domazet-Loso, 2011; Neme & Tautz, 2013). Multiple studies have shown that young genes (i.e. those genes whose recognised homologs are only present in closely related species (Domazet-Loso et al. 2007) evolve faster than old genes (Thornton & Long, 2002; Domazet-Loso & Tautz, 2003; Krylov et al. 2003; Daubin & Ochman, 2004; Alba & Castresena, 2005; Wang et al. 2005; Cai et al. 2006; Wolf et al. 2009; Cai & Petrov, 2010; Zhang et al. 2010; Vishnoi et al. 2010; Tautz & Domazet-Loso, 2011; Cui et al. 2015). Cai and Petrov (2010) found clear evidence for the role of non-adaptive evolution in this relationship but no evidence for adaptive evolution. However, there is an expectation that young genes will be further from their evolutionary optimum than old genes, and hence that they should undergo rapid adaptive evolution when they are born. There is some limited evidence for this; the *jingwei* gene, which appeared very recently in the *Drosophila* phylogeny is evolving very rapidly, with 80% of the amino acid substitutions estimated to have been due to adaptive evolution (Long & Langley, 1993).

Recombination is expected to affect the probability that both advantageous and deleterious mutations are fixed, due to its ability to reduce Hill-Robertson interference between selected mutations (Hill & Robertson 1966; Marais & Charlesworth, 2003). Rates of adaptation have been shown to be strongly positively correlated to recombination rate in *Drosophila* (Presgraves, 2005; Betancourt et al. 2009; Arguello et al. 2010; Mackay et al 2012; Campos et al. 2014; Castellano et al. 2016; Moutinho et al. 2019) and *Arabidopsis* (Moutinho et al. 2019), and rates of non-adaptive evolution to be negatively correlated in both *Drosophila* and *Arabidopsis* species (Moutinho et al. 2019).

In summary, a number of factors have been shown to correlate to rates of protein evolution, and in some of these cases the relative roles of adaptive and non-adaptive evolution have been disentangled. However, relatively little work has been done on these questions in hominids. We addressed these questions by considering the role of gene age, RR, gene expression, protein length and gene function in determining rates of both adaptive and non-adaptive evolution. To disentangle the effects of adaptive and non-adaptive evolution we use an extension of the McDonald-Kreitman test which estimates these quantities taking into account the distribution fitness effects of new mutations.

## Results

We set out to investigate whether a number of gene-level factors affect the rate of adaptive and non-adaptive evolution in hominids – the rate of recombination (RR), gene age, the level of gene expression, gene length and gene function. We measure the rates of adaptive and non-adaptive evolution using the statistics *ω*_*a*_ and *ω*_*na*_, which are estimates of the rate of evolution relative to the mutation rate. We estimated both statistics using an extension of the McDonald-Kreitman method, in which the pattern of substitution and polymorphism at neutral and selected sites is used to infer the rates of substitution, taking into account the influence of slightly deleterious mutations. We use the method implemented in Grapes (Galtier, 2016), which is a maximum likelihood implementation of the second method proposed by Eyre-Walker and Keightley (2009). Note that genes are grouped together according to the factors analysed, since most genes have relatively little polymorphism data, and this makes estimating the rate of adaptive evolution for individual genes impractical.

We estimated *ω*_*a*_ and *ω*_*na*_ using 16,344 genes for the divergence between humans and chimpanzees using African SNPs from the 1000 genomes data (The 1000 Genomes Project Consortium, 2015). We find that the average rate of adaptive evolution is approximately five-fold lower than the rate of non-adaptive evolution (*ω*_*a*_ = 0.037 (95% confidence intervals estimates using bootstrapping 0.035 and 0.039) versus *ω*_*na*_ = 0.192 (0.190,0.194)). The proportion of substitutions that are adaptive, α, is estimated to be 0.162, which is close to previous recent estimates (Eyre-Walker & Keightley, 2009; Boyko et al. 2008; Messer & Petrov 2013).

### Adaptive evolution

The rate of adaptation is expected to be retarded in regions of low recombination because of Hill-Robertson interference, and we do indeed find that the rate of adaptive evolution is significantly positively correlated to the rate of recombination in hominids (Figure 1a; (r=0.737, p<0.001)). A similar positive correlation has previously been observed in *Drosophila* (Presgraves, 2005; Betancourt et al. 2009; Arguello et al. 2010; Mackay et al. 2012; Campos et al. 2014; Castellano et al. 2016). In the most detailed study of this relationship in *Drosophila*, Castellano et al. (2016) found that the rate of adaptive evolution increases with RR, but that it asymptotes, suggesting that above a certain level of recombination, Hill-Robertson interference has little effect. However, we do not observe an asymptote in hominids (figure 1a). Since there is a large difference in average recombination between the two groups with the highest recombination rate, we repeated the analysis with 50 recombination bins; although we still observe a significant positive correlation between *ω*_*a*_ and RR (r=0.582, p<0.001), the analysis failed to reveal any signal of an asymptote (figure S1).

**Figure 1:**
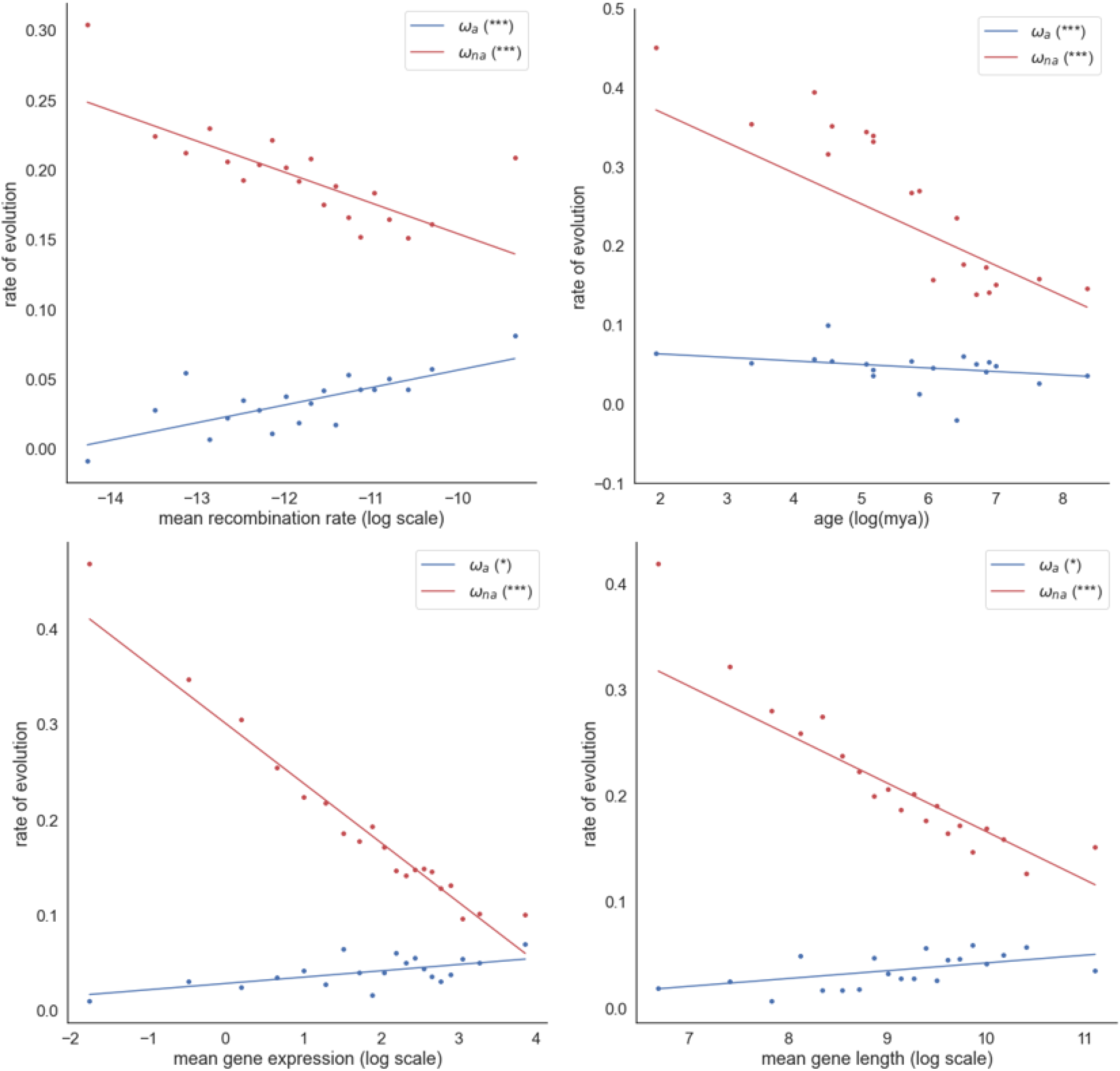
Estimates of *ω*_*a*_ and *ω*_*na*_ plotted against the mean recombination rate (a), gene age (b), mean gene expression (c) and mean protein length (d). The respective significance of each correlation is shown in the plot legend, (*P < 0.05; **P < 0.01; ***P < 0.001; “.” 0.05 ≤ P < 0.10) for *ω*_*a*_ and *ω*_*na*_). Also shown is the line of best fit through the data. An unweighted regression is fitted to the estimates of *ω*_*a*_ and *ω*_*na*_.

Young genes have been shown to evolve faster than old genes (Thornton & Long, 2002; Domazet-Loso & Tautz, 2003; Krylov et al. 2003; Daubin & Ochman, 2004; Alba & Castresena, 2005; Wang et al. 2005; Cai et al. 2006; Wolf et al. 2009; Cai & Petrov, 2010; Zhang et al. 2010; Vishnoi et al. 2010; Tautz & Domazet-Loso, 2011; Cui et al. 2015). There is an expectation that young genes will undergo faster rates of adaptive evolution because they are further from their adaptive optima (Wright, 1931, 1932), and we do indeed find a significant negative correlation between *ω*_*a*_ and gene age (r=-0.404, p=0.012) in hominids (figure 1b).

Highly expressed genes have been shown to exhibit lower rates of protein evolution in both eukaryotes (Pal et al. 2001; Subramanian & Kumar, 2004; Wright et al. 2004; Lemos et al. 2005) and prokaryotes (Rocha & Danchin, 2004). Moutinho, et al. (2019) found significant negative correlations in *Drosophila* species between *ω*_*a*_ and both gene expression and protein length. Intriguingly, the correlations are reversed in hominids, with both correlations being significantly positive (gene expression: r=0.642, p=0.002; protein length: r=0.597, p=0.005) (figures 1c & 1d).

### Independent effects

Our measure of adaptive evolution, *ω*_*a*_, is significantly positively correlated to RR, expression and protein length, and negatively to gene age. However, the rate of recombination, gene age, gene expression and protein length are all significantly, or marginally significantly, correlated to each other (Table 1) so it is important to determine whether each factor has an independent effect on the rate of adaptive evolution; i.e. the correlation between Y and X, might be due to the fact that each is correlated to a third factor Z, and with no variation in Z there is no correlation between Y and X. To investigate this, we conducted two analyses. In the first instance, we repeated our analyses controlling for each factor in turn by taking the values of the co-correlate around the modal value. We took the modal value and 0.5 standard deviations either side; this significantly reduced the standard deviation of the co-correlate within each analysis, largely controlling for this factor (Table 1). However, controlling for each factor this way reduces the data set considerably, so we also ran an analysis in which we calculated the expected correlation between two variables under the assumption that they are correlated solely because of their correlation to a third variable. It can be shown that if the correlation between Y and Z is *r*_*YZ*_ and that between X and Z is *r*_*XZ*_, then expected correlation between Y and X due to the covariation with Z is 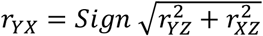, where Sign is positive if both *r*_*YZ*_ and *r*_*XZ*_ are positive or negative, and negative otherwise. In both analyses, we investigate factors that could generate an artefactual correlation – e.g. the correlation between Y and X cannot be due to covariation with Z, if Y and X are positively correlated, but the correlation between Y and Z is positive but the correlation between X and Z is negative.

**Table 1:** The correlation between the gene age, gene expression, gene length and recombination rate; logs were taken of all variables. The CV column is the coefficient of variation of the factor for all the data. The final column is the CV of the restricted data (i.e. when we control for the factor in question by restricting the analysis to genes with the modal value +/-0.5 standard deviations).

|  | gene<br>expression | gene length | recombination<br>rate | CV | CV of near<br>modal values |
| --- | --- | --- | --- | --- | --- |
| gene age | 0.868 (***) | 0.860 (***) | -0.621 (**) | 1.385 | 0.381 |
| gene<br>expression |  | 0.437 (***) | -0.035 (***) | 1.451 | 0.411 |
| gene length |  |  | 0.101 (***) | 1.727 | 0.496 |
| recombination<br>rate |  |  |  | 1.143 | 0.325 |

Our two analyses suggest that there is a direct association between *ω*_*a*_ and RR; when we control for age and length, we find that although the correlation is no longer significant when we control for either variable, the correlation does remain positive, and the observed correlations are significantly greater than the predicted correlation (Table 2). The analysis also suggests that there is a direct association between *ω*_*a*_ and age, because the correlation remains significantly negative when we control for RR, and the predicted correlation is significantly smaller in magnitude than the observed correlation. However, the results with gene expression and length are less clear; when each variable is controlled for in the analysis of the other, the correlation becomes non-significant (Table 2). The observed correlation between *ω*_*a*_ and expression is significantly greater than the predicted correlation, using length as the covariate, whereas the opposite is not true; this would seem to suggest that there is a direct correlation between *ω*_*a*_ and expression, and that the correlation between *ω*_*a*_ and length may be due to the fact that both are correlated to expression. However, the evidence is not strong in support of this hypothesis.

**Table 2.** The observed correlation between Y and X controlling for a covariate, Z, and the observed and predicted correlation between Y and X assuming the relationship is solely due to the correlation between each variable and a third factor Z. The final column gives the proportion of 100 botstrap replicates in which the predicted correlation divided by the observed correlation is greater than one – i.e. the predicted correlation is larger in magnitude.

| Y variate | X variate | Observed r | Z variate | Observed r - controlling for Z | Predicted r | Predicted/observed > 1 |
| --- | --- | --- | --- | --- | --- | --- |
| $\omega_a$ | RR | 0.74*** | Age | 0.25 | 0.15 | 0 |
| $\omega_a$ | RR | 0.74*** | Length | 0.43 | 0.086 | 0 |
| $\omega_a$ | Age | -0.40* | RR | -0.58* | -0.093 | 0.02 |
| $\omega_a$ | Expression | 0.64** | Length | 0.00 | 0.38 | 0.03 |
| $\omega_a$ | Length | 0.60** | RR | 0.64** | 0.091 | 0 |
| $\omega_a$ | Length | 0.60** | Expression | 0.25 | 0.37 | 0.13 |
| $\omega_{na}$ | RR | -0.73*** | Length | -0.54* | -0.34 | 0 |
| $\omega_{na}$ | Age | -0.91*** | Expression | -0.76** | -0.76 | 0 |
| $\omega_{na}$ | Age | -0.91*** | Length | -0.87*** | -0.75 | 0 |
| $\omega_{na}$ | Expression | -0.98*** | Age | -0.74*** | -0.90 | 0 |
| $\omega_{na}$ | Expression | -0.98*** | Length | -0.61** | -0.95 | 0.01 |
| $\omega_{na}$ | Length | -0.94*** | RR | -0.91*** | -0.42 | 0 |
| $\omega_{na}$ | Length | -0.94*** | Age | -0.49* | -0.88 | 0 |
| $\omega_{na}$ | Length | -0.94*** | Expression | -0.71*** | -0.89 | 0 |

There is another factor that needs to be controlled for in any analysis of age -fast evolving genes are harder to identify in more distant species, and this can lead to an artefactual correlation between the age of a gene and the rate of evolution. The distribution of non-synonymous substitution rates is bimodal, with many genes having *d*_*N*_ = 0. We took genes around the second mode, those with rates between 0.002 and 0.007. This reduces our dataset from 15,439 to 4,961 genes, and as a consequence we had to combine multiple age categories together. We find no significant correlation between *ω*_*a*_ and age when we do this (r=0.413, p=0.270), suggesting that the correlation between *ω*_*a*_ and age might be an artifact of the problems in identifying fast evolving genes in older taxa.

### Controlling for biased gene conversion

Biased gene conversion (BGC) can potentially impact estimates of the rate of adaptive evolution, either by increasing the fixation probability of S over W neutral alleles (Galtier & Duret, 2007; Berglund et al. 2009; Ratnakumar et al. 2010; Rousselle et al. 2020), or by promoting the fixation of slightly deleterious S alleles (Duret & Galtier, 2009; Glemin, 2010; Necsulea et al. 2011; Lachance & Tishkoff, 2014; Rousselle et al. 2019). To investigate whether BGC affects our results we can leverage some of the results above. The correlation between *ω*_*a*_ and either age and protein length remains significant if we control for RR (Table 2) (supplementary figures, S3a and S6a respectively), so it seems that BGC is unlikely to be responsible for these correlations. If we control for RR in the regression between *ω*_*a*_ and expression, we find that the correlation remains, suggesting that this correlation is also not due to BGC (r=0.780, p<0.001) (supplementary figure S5a).

To investigate whether the correlation between *ω*_*a*_ and RR is due to BGC we performed a different analysis restricting the analysis to those polymorphisms and substitutions that are unaffected by BGC – i.e. A<>T and G<>C changes. This reduces our dataset to about 20% of its previous size. We find that there is still a positive correlation, although this is only marginally significant (r=0.102, p=0.093) (Supplementary figure, S2).

In conclusion, *ω*_*a*_ is positively correlated to RR, protein length and gene expression level, and to a large extent these correlations survive controlling for each other and BGC; the exceptions are protein length when expression is controlled for, and the positive relationship between *ω*_*a*_ and RR when BGC is controlled for; this latter correlation remains marginally significant.

### Non-adaptive evolution

We repeated the analysis above for the rate of non-adaptive evolution. We find that *ω*_*na*_ is highly significantly negatively correlated to RR, gene age, length and expression (Table 2; Figure 1). All of these correlations remain significant when controlling for potentially confounding factors, and the observed correlation is significantly greater in magnitude than the predicted correlation (Table 2). Hence, we can conclude that all four factors have significant independent effects on *ω*_*na*_. As with the analysis of *ω*_*a*_ it is possible that these correlations are due to BGC. However, if we control for RR in our analyses we find that all the negative correlations persist (gene age: r=-0.886, p<0.001; gene length: r=-0.910 p<0.001; gene expression: r=0.989, p<0.001). In the case of the correlation between *ω*_*na*_ and RR, if we restrict the analysis to G<>C and A<>T mutations we find that *ω*_*na*_ remains significantly positively correlated to RR (r=-0.648, p<0.001).

### Gene function

In the second part of our analysis, we consider the effect of gene function on the rate of adaptive and non-adaptive evolution. It has previously been demonstrated that genes whose products interact with viruses – viral interacting proteins (VIPs) – have higher rates of adaptive evolution than other genes in primates (Enard et al. 2016). We confirm this pattern. In our analysis, in which we have used a different method and statistic to estimate the rate of adaptive evolution, we find that the rate of adaptive evolution amongst VIPs is approximately 40% greater than in non-VIPs (*ω*_*a*_ = 0.052 versus 0.032), a difference that is highly significant (p<0.001). This pattern is consistent across almost all GO categories that have at least 100 genes, supporting the results of Enard et al. (2016) (figure 2).

**Figure 2:**
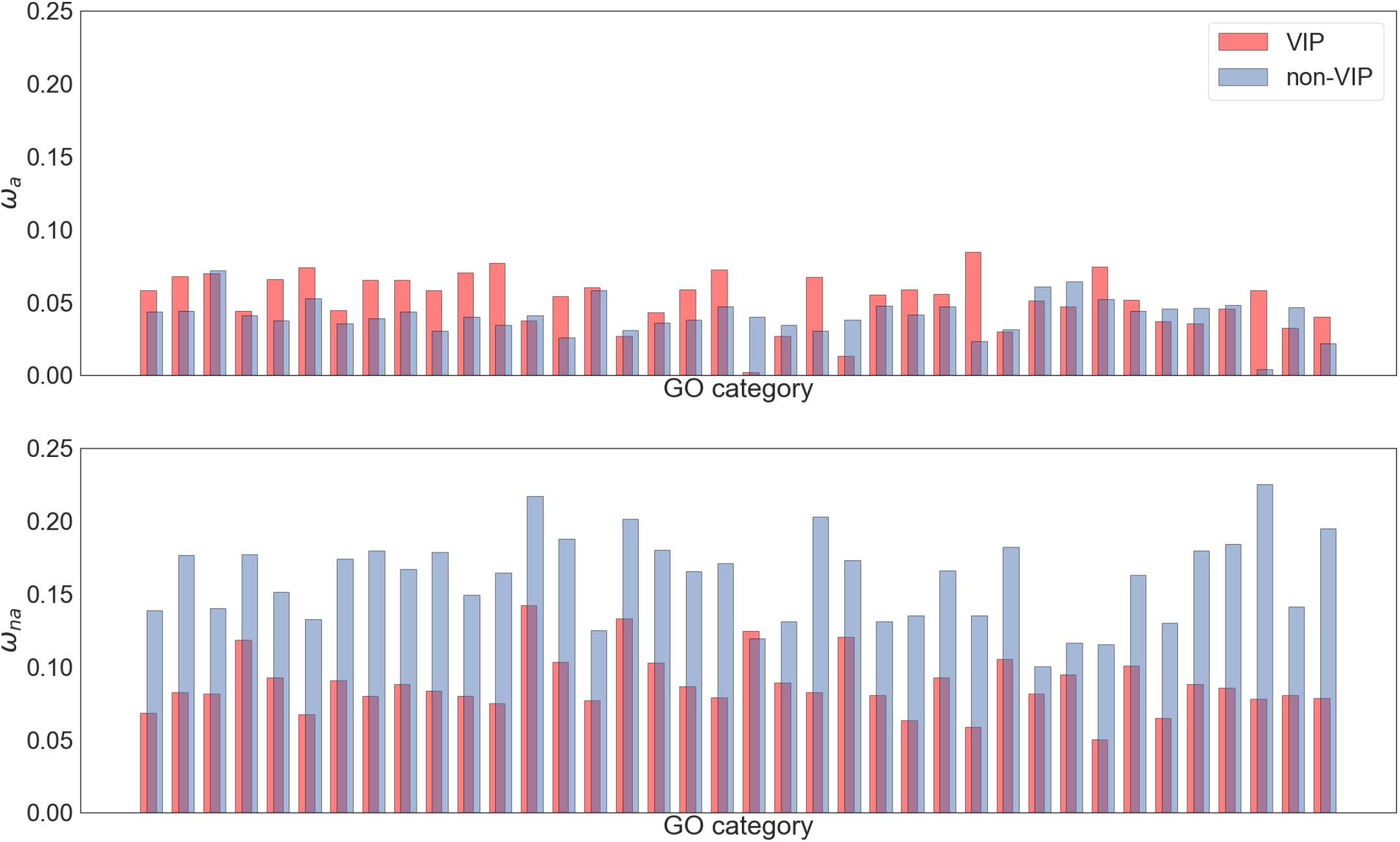
Estimates of *ω*_*a*_ (top) and *ω*_*na*_ (bottom) for GO categories that contain >100 viable VIP and non-VIP genes.

It is evident however, that there is substantial variation between GO categories for non-VIP genes, and this variation is significant, taking into account that individual genes can contribute to multiple GO categories (p=0.0012). This pattern is replicated if we include GO categories which do not include VIP proteins (p=0.0010). The GO categories which have the highest rate of adaptive evolution are ubiquitin protein ligase binding, and protein kinase binding (table 3).

**Table 3:** Top 10 GO categories, ranked by rate of adaptive substitution.

| GO category | $\omega_a$ | $\omega_a$ 95% confidence intervals |
| --- | --- | --- |
| ubiquitin protein ligase binding | 0.0843 | 0.0702 - 0.0995 |
| protein kinase binding | 0.0804 | 0.0698 - 0.0914 |
| sequence-specific DNA binding | 0.0735 | 0.0633 - 0.0842 |
| DNA-binding transcription factor activity | 0.0719 | 0.0628 - 0.0812 |
| transcription factor complex | 0.0682 | 0.0496 - 0.0883 |
| transcription by RNA polymerase II | 0.0673 | 0.0518 - 0.0836 |
| negative regulation of apoptotic process | 0.0671 | 0.0552 - 0.0796 |
| chromatin organization | 0.0669 | 0.0567 - 0.0775 |
| DNA-binding transcription activator activity | 0.0649 | 0.0524 - 0.078 |
| transcription coactivator activity | 0.0648 | 0.0519 - 0.0786 |

What are the relative contributions of GO category and VIP status to the variation in the rate of adaptive evolution – i.e. is most of the variation in the rate of adaptive evolution due to whether the gene encodes a VIP or not, or is most of the variation due to other functional considerations? To investigate this, we performed a two-way analysis of variance on *ω*_*a*_ and estimated the variance components. We find that the distinction between VIP and non-VIP contributes approximately 5x the variance in *ω*_*a*_ as the variation between GO categories, suggesting that whether a gene encodes a VIP has a major effect on its rate of adaptation (supplementary table, S1).

But what of non-adaptive evolution? If we divide our data into genes that interact with viruses and those that do not, we find that rates of non-adaptive evolution are substantially higher in non-VIP genes (*ω*_*na*_ = 0.198 vs 0.101). As Enard et al. (2016) found, this pattern is replicated across GO categories (Figure 2). There is substantial and significant variation in *ω*_*na*_ across GO categories excluding VIP genes, taking into account that individual genes can contribute to multiple GO categories (p<0.001). This pattern is replicated if we include GO categories which do not include VIP proteins (p<0.001). The GO categories that have the highest non-VIP rates of non-adaptive evolution are both related to immune system response (table 4). If we partition the variance between VIP/non-VIP and GO categories we find that the distinction between VIP and non-VIP contributes over 8x the variance in *ω*_*na*_ as the variation between GO categories, suggesting that whether a gene encodes a VIP has a major effect on its rate of non-adaptive evolution (supplementary table, S2) as well as its rate of adaptation.

**Table 4:**
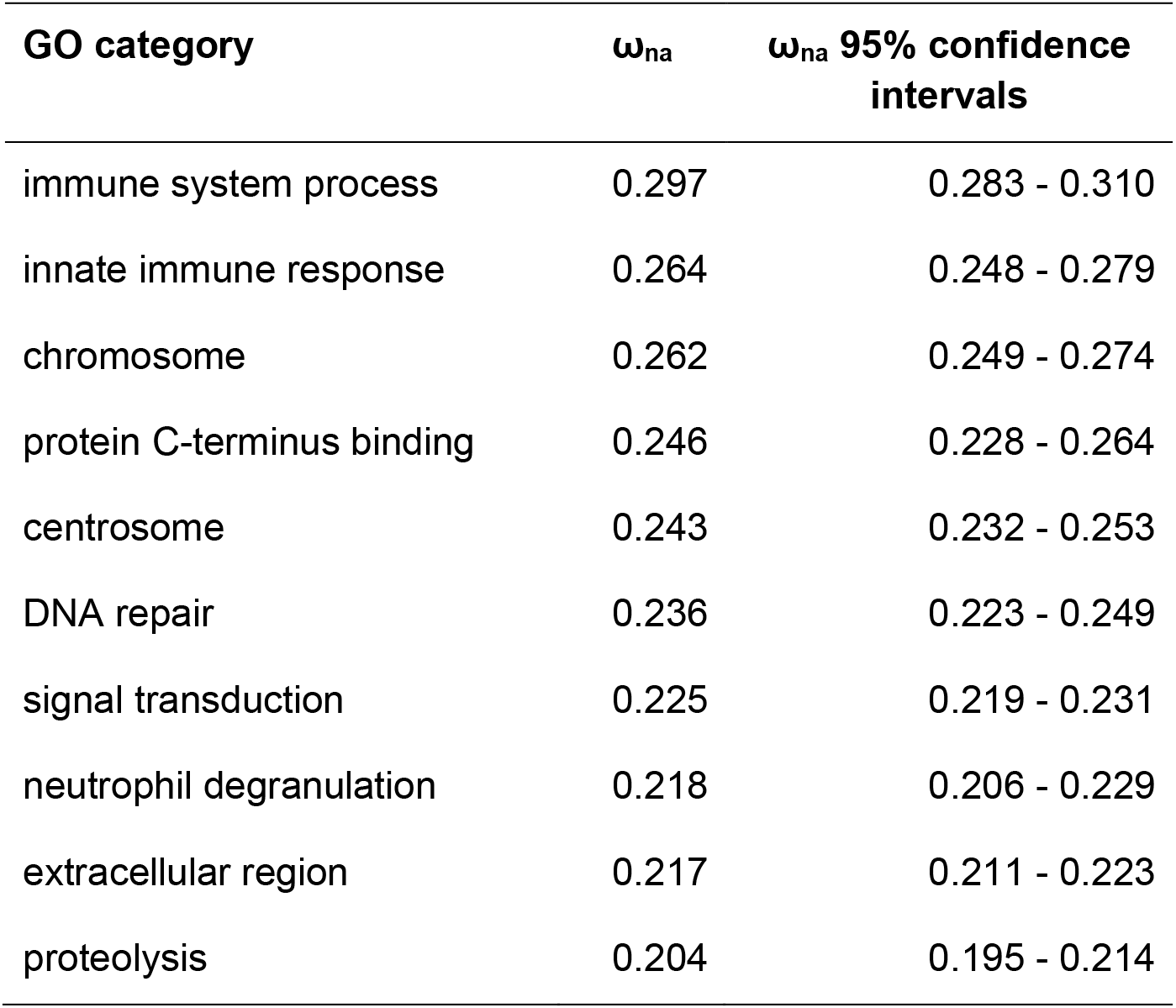
Top 10 GO categories, ranked by rate of non-adaptive substitution

| GO category | $\omega_{na}$ | $\omega_{na}$ 95% confidence intervals |
| --- | --- | --- |
| immune system process | 0.297 | 0.283 - 0.310 |
| innate immune response | 0.264 | 0.248 - 0.279 |
| chromosome | 0.262 | 0.249 - 0.274 |
| protein C-terminus binding | 0.246 | 0.228 - 0.264 |
| centrosome | 0.243 | 0.232 - 0.253 |
| DNA repair | 0.236 | 0.223 - 0.249 |
| signal transduction | 0.225 | 0.219 - 0.231 |
| neutrophil degranulation | 0.218 | 0.206 - 0.229 |
| extracellular region | 0.217 | 0.211 - 0.223 |
| proteolysis | 0.204 | 0.195 - 0.214 |

## Discussion

It has been previously shown that the rate of evolution correlates to a number of factors including RR (Presgraves, 2005; Betancourt et al. 2009; Arguello et al. 2010; Mackay et al 2012; Campos et al. 2014; Castellano et al. 2016; Moutinho et al. 2019), gene age (Thornton & Long, 2002; Domazet-Loso & Tautz, 2003; Krylov et al. 2003; Daubin & Ochman, 2004; Alba & Castresena, 2005; Wang, et al., 2005; Cai, et al., 2006; Wolf, et al., 2009; Cai & Petrov, 2010; Zhang et al. 2010; Vishnoi et al. 2010; Tautz & Domazet-Loso, 2011; Cui, et al., 2015), expression level (Pal et al. 2001; Rocha & Danchin, 2004; Subramanian & Kumar, 2004; Wright et al. 2004; Lemos et al. 2005; Moutinho et al. 2019) and protein length (Zhang, 2000; Lipman et al. 2002; Liao et al. 2006; Moutinho et al. 2019). In addition, the rate of evolution has been shown to vary with gene function (Clark et al. 2003; Nielsen et al. 2005; Chimpanzee Sequencing and Analysis Consortium, 2005). In this study we have correlated each of these factors to *ω*_*a*_ and *ω*_*na*_ in hominids, allowing us to disentangle the effects of adaptive and non-adaptive evolution. We find that *ω*_*a*_ is correlated to all four factors, and that when we control for each factor in turn, there is evidence for an independent influence of RR, gene age and gene expression. These correlations remain when controlling for the effects of biased gene conversion, although the relationship with RR is only marginally significant. However, the correlation with gene age could be an artefact of fast evolving genes having higher rates of adaptive evolution and being more difficult to identify in older taxa; when we control for the rate at which a protein evolves the negative correlation between *ω*_*a*_ and gene age becomes non-significant suggesting that this pattern might be an artefact.

In contrast, we find that all four factors have significant independent effects on *ω*_*na*_, and that all of these remain significant when we control for each in turn, and control for BGC. Several studies on both eukaryotes (Pal et al. 2001; Subramanian & Kumar 2004; Wright et al. 2004; Lemos et al. 2005; Moutinho et al. 2019) and prokaryotes (Rocha & Danchin 2004) have demonstrated that more highly expressed genes have lower rates of protein sequence evolution. Our results support these previous findings, with the negative correlation between *ω*_*na*_ and gene expression suggesting that more highly expressed genes are under greater constraint in hominids. Drummond et al. (2005) suggest a general hypothesis that more highly expressed genes evolve slowly (i.e. are under higher selective constraint) because of the selection against the expression level cost of protein misfolding, wherein selection acts by favoring protein sequences that accumulate less translational missense errors. We also find a significant negative correlation between *ω*_*na*_ and gene length. This supports former studies that have shown that smaller genes evolve more rapidly (Zhang 2000; Lipman et al. 2002; Liao et al. 2006; Moutinho et al. 2019), suggesting that smaller protein-coding regions are under more relaxed purifying selection.

### Gene function analyses

Our analyses of VIP and non-VIP genes show that a high proportion of the variance in protein evolution in hominids is accounted for by whether or not a gene interacts with viruses, a result that corroborates Enard et al.’s (2016) findings. By disentangling the rates of adaptive and non-adaptive evolution, we find that VIP genes are under less constraint than non-VIPs, and that VIPs exhibit a higher rate of adaptive evolution. We also estimate the variance components using two-way analyses of variance, finding that the distinction between VIP and non-VIP contributes about 5x the variance in *ω*_*a*_, and 8x the variance in *ω*_*na*_ as the variation between GO categories, suggesting that whether a gene encodes a VIP has a major effect on its rate of adaptation and non-adaptation (supplementary table, S1). These results could explain why there appears to be little variation in the rate of adaptive evolution across biological functions categorised using Gene Ontology (Bierne & Eyre-Walker, 2004), with viruses acting across a range of biological functions likely to be a key factor in these estimates.

Our study is likely to underestimate the amount of adaptive evolution attributable to viruses, for reasons outlined by Enard et al (2016). Briefly, we used the categorisation of VIPs and non-VIPs provided by Enard et al (2016). However new VIPs are being discovered regularly, suggesting there are some VIPs that were not included in our analysis. Secondly, the categorisation of VIP and non-VIP necessarily cannot account for proteins that adapt to viruses but do not physically interact with them (e.g. in proteins that are upstream or downstream of VIPs in signaling cascades).

### No asymptote in the correlation between ω_a_ and RR

Both Campos et al. (2014) and Castellano et al. (2016) found that there is a positive relationship between the rate of adaptive evolution and RR in *Drosophila*. However, Castellano et al. (2016) showed that the positive correlation between RR and *ω*_*a*_ asymptotes in *Drosophila*, suggesting that above a certain level of recombination Hill-Robertson interference has little effect. In this study we find no evidence of this asymptote in hominids for either the rate of adaptive or non-adaptive evolution, suggesting that most coding sequences may experience some level of HRi. This is perhaps not unexpected. The level of HRi will depend on several factors -the effectiveness of recombination in breaking down associations, the density of selected sites and the mutation rate to alleles that are subject to selection; if weakly selected mutations are responsible for HRi then the effective population size and the level of nearly neutral genetic diversity will also be important. Recombination is a considerably more effective force in Drosophila; linkage disequilibrium (LD) decays over a scale of 10s of base pairs (Mackay et al. 2012) rather than the 10,000s that we observe in humans (The 1000 Genomes Project Consortium, 2015). This 1000-fold difference in the effectiveness of recombination is likely to more than compensate for the fact that humans have ∼20-fold greater genome size, and a higher rate of deleterious mutation (2.1 in humans (Lesecque et al. 2012) to 1.2 in Drosophila (Haag-Liautard et al. 2007) respectively).

### Gene age

Cai and Petrov (2010) found that older genes exhibit a lower rate of protein evolution (as measured by the Ka/Ks ratio) than younger genes. The authors demonstrated that this was at least in part due to stronger purifying selection acting on older genes than on younger ones, by showing that levels of non-synonymous to synonymous polymorphism were lower in older genes. Our findings corroborate these results, with the strong negative correlation between *ω*_*na*_ and gene age showing that older genes are under a lower rate of protein evolution than younger genes. However, we also find a significant negative correlation between gene age and the rate of adaptive evolution, *ω*_*a*_, whilst Cai and Petrov found no such correlation. There are two potential causes of this discrepancy. Firstly, for this analysis Cai and Petrov group genes by their age based on lineage specificity (LS), that is, how specifically a gene and orthologs of a gene are distributed on a given phylogeny (Cai et al. 2006), whilst we group our genes by phylostratigraphic category (PL), that is, where genes are ranked by phylostratigraphic category based on their earliest ortholog (Domazet-Loso et al. 2007). Each method has its limitations. Because the LS method relies on the phylogenetic profiles of individual genes, Cai and Petrov removed genes with patchy distributions (Cai et al. 2006), resulting in 10,032 of 20,150 genes being removed from the dataset for having irregular phylogenetic profiles. The PL method relies on parsimony and assumes that a gene family can be lost, but cannot re-evolve in different lineages (Domazet-Loso et al. 2007), meaning that those genes that would be removed using the LS method are maintained in the PL method. By using the PL method, our dataset contained 15,439 grouped into 19 phylostratigraphic bins. Secondly, Cai and Petrov obtained divergence and polymorphism data from the compiled Applera dataset (Bustamante et al. 2005; Lohmueller, et al., 2008) of 39 humans (19 African Americans and 20 European Americans), whilst we have used data from the 661 African samples within the 1000 genomes dataset (The 1000 Genomes Project Consortium, 2015). Notably, the African population has undergone a more stable demographic history than Europeans, who carry proportionally more deleterious genetic variation, which Lohmueller et al (2008) ascribe to the bottleneck encountered by the Eurasian population at the time of the migration out of Africa. This higher proportion of segregating deleterious alleles will inevitably affect estimates of the rate of adaptive evolution, but not the ratio of non-synonymous and synonymous substitution rates (the latter of which yields a strong correlation with gene age using both the PL and LS methods in Cai and Petrov’s study).

### The effect of population contraction

It has been shown previously that the MK test can generate artifactual evidence of adaptive evolution if some nonsynonymous mutations are slightly deleterious and the population in question has undergone recent expansion, because selection is more effective during the polymorphism phase than during the divergence phase (McDonald & Kreitman, 1991; Eyre-Walker, 2002). Although, the effective population size in humans has increased recently, the effective population size is considerably reduced from that in the human-chimpanzee ancestor (Hobolth et al. 2007; Burgess and Yang 2008; Prado-Martinez et al. 2013; Schrago, 2014). This population contraction can depress the signal of adaptive evolution in humans. Furthermore, we will show elsewhere that if a factor, for example gene age, is correlated to the mean strength of selection against deleterious mutations, population size change will generate an artifactual correlation between that factor and the rate of adaptive evolution. The direction of this correlation depends on the direction of the correlation between the mean strength of selection acting against deleterious mutations and the factor in question and whether the population has expanded or contracted; for example, if factor X is positively correlated to the mean strength of selection (i.e. selection is stronger against genes with larger values of X), then population contraction will induce an artifactual positive correlation between *ω*_*a*_ and X.

Figure 3 shows that all four factors are positively correlated to the log mean strength of selection against deleterious mutations, estimated from the site frequency spectrum (gene age: r=0.916, p<0.001; RR: r=0.828, p<0.001; gene length: r=0.818, p<0.001; gene expression: r=0.948, p<0.001). Population contraction undergone by humans should therefore tend to induce a positive correlation between *ω*_*a*_ and each factor in our analysis. This artifactual positive correlation is contrary to the negative correlation that we observe between *ω*_*a*_ and age (Figure 1). This may be one reason why we observe a weaker correlation between gene age and the rate of adaptive evolution in hominids compared with *Drosophila* and *Arabidopsis* species (Moutinho et al. unpublished). However, population contraction might also be responsible for the positive correlation between *ω*_*a*_, RR, protein length and expression. Because *ωna* is estimated exclusively from polymorphism phase data, we do not expect the correlations between *ωna* and our four factors to be affected by the population contraction.

**Figure 3:**
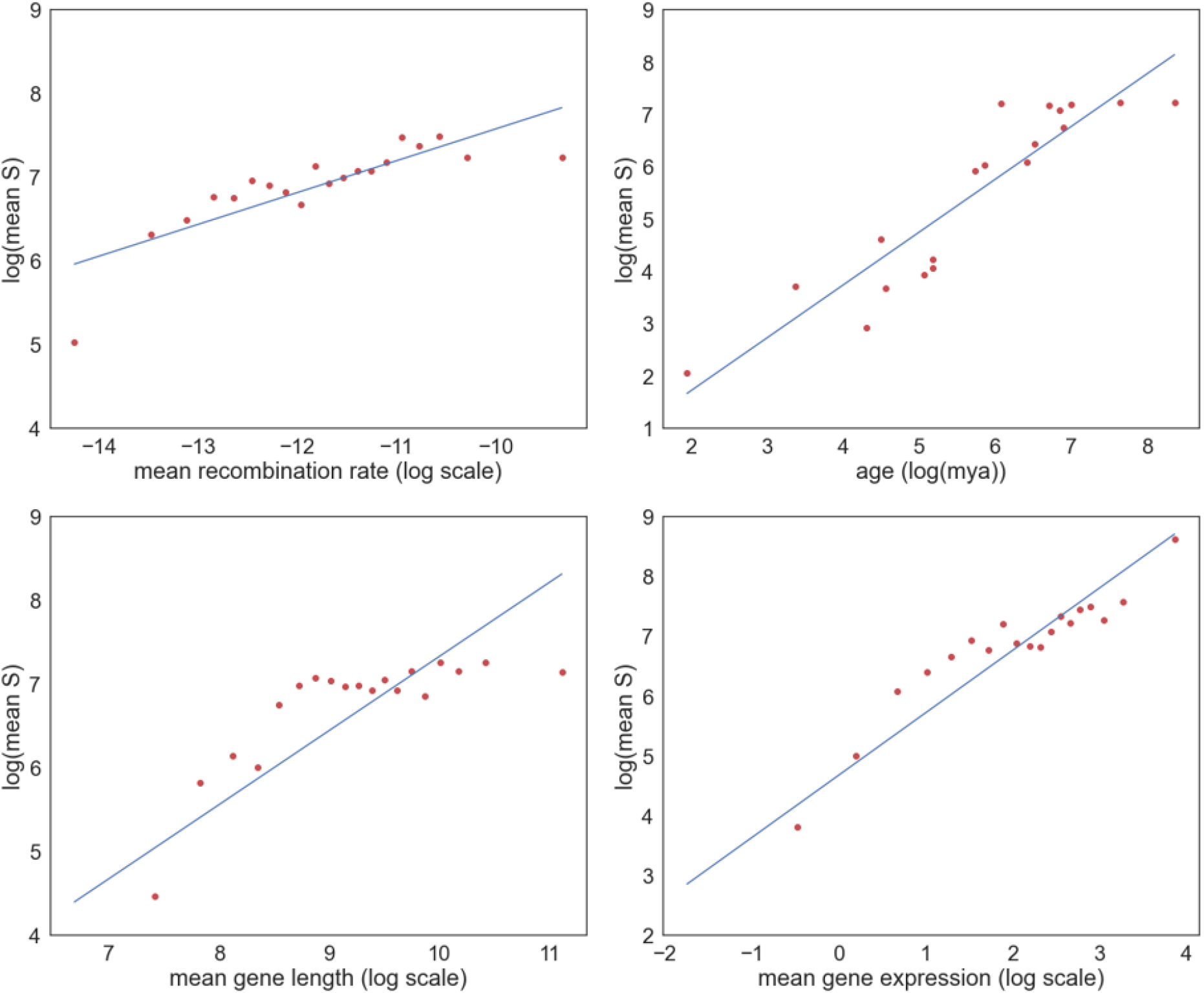
Correlation between the log of the mean strength of selection against deleterious mutations and gene age (top left), RR (top right), gene length (bottom left), gene expression (bottom right). A linear regression has been fitted to each dataset.

In summary, we observe a significant correlation between the rate of adaptive evolution, RR, protein length and gene expression, and a negative correlation between the rate of adaptive evolution and gene age. However, we cannot be very confident that any of these correlations are genuine; the positive correlation between *ω*_*a*_, RR, protein length and gene expression might be due to an artifact of population size contraction, and the correlation between *ω*_*a*_ and age might be due to the problems of identifying rapidly evolving genes, with high values of *ω*_*a*_, in more distant taxa. In contrast, the rate of non-adaptive evolution is independently negatively correlated to all factors. We have confirmed that whether a protein interacts with viruses is an important factor in determining whether a gene undergoes high rates of adaptive and non-adaptive evolution, however we also demonstrate that there is significant variation between GO categrories, even when this factor is controlled for.

## Materials and methods

### Data

We obtained orthologous human and chimpanzee gene sequences from the Ensembl biomart (Yates et al. 2019) for the human GRCh38 and Pan_tro_3.0 genome builds. We aligned these orthologs using MUSCLE (Edgar, 2004). After filtering out genes with gaps that were not a multiple of 3 we were left with 16,344 pairwise alignments. Proportions of synonymous and non-synonymous substitutions were estimated using codeml from the PAML package (Yang, 2007) program. We used polymorphism data from the African superpopulation of the 1000 genomes dataset (The 1000 Genomes Consortium, 2015) to construct our site frequency spectra, with rates of adaptive (*ω*_*a*_) and non-adaptive (*ω*_*na*_) evolution estimated using Grapes (Galtier, 2016), under the “GammaZero” model. We used African SNPs because the African population has been subject to relatively simple demographic processes (Gravel et al. 2011). Confidence intervals on our estimates of *ω*_*a*_ and *ω*_*na*_ were generated by bootstrapping the dataset by gene.

Gene ages were obtained from Litman and Stein (2019). In this dataset genes are ranked by phylostratigraphic category based on their earliest ortholog. Gene lengths were obtained by taking the total coding sequence length of the longest transcript of each protein, whilst gene expression data was obtained from the Expression Atlas database (Papatheodorou et al. 2019), wherein the baseline experiment E-MTAB-5214 was used. This data is from the GTEx genotype-tissue expression analysis of 53 tissue samples (GTEx Consortium, 2015). We estimated the arithmetic mean expression value across tissues for each gene, and binned gene by mean gene expression of 20 roughly equally sized bins (each containing 808-811 genes). Recombination rate maps were obtained from Spence and Song (2019), and the mean recombination rate was calculated between the start and end of the largest transcript for each gene. GO category information was obtained from Ensembl’s Biomart (Ashburner et al. 2000; The Gene Ontology Consortium, 2021; Yates et al. 2019).

### Correlating factors with rates of adaptive and non-adaptive evolution

To correlate the rates of adaptive and non-adaptive evolution with each of recombination rate, protein length and gene expression we binned our genes into 20 roughly equal sized bins. For gene age we binned data by phylostratigraphic category, of which there were 19. To control for biased gene conversion in our recombination rate analysis we restricted the analysis to those polymorphisms and substitutions that are unaffected by biased gene conversion – i.e. A<>T and G<>C changes. This reduced our dataset to about 20% of its previous size.

To investigate whether factors were independently correlated to *ω*_*a*_ and *ω*_*na*_ we ran the analysis controlling for each of the other three factors in turn. We controlled for each factor by taking the values of the co-correlate close to the modal value. We took the modal value and 0.5 standard deviations either side which reduces the standard deviation of the co-correlate within each analysis. Because this reduces the data set considerably, we also ran an analysis in which we predicted the correlation coefficient between Y and X under the assumption that they are only correlated to each other because they are both correlated to Z. If *r*_*YZ*_ is the correlation between Y and Z, then *r*_YZ_ ^*2*^ is the proportion of variance in Y explained by Z, and vice versa. Hence, the proportion of variance explained in Y by X, because of their mutual correlation to Z is *r* ^*2*^ *r* ^*2*^. Hence the expected correlation coefficient between Y and X is 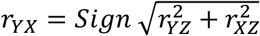, where Sign is positive if both *r*_*YZ*_ and *r*_*XZ*_ are positive or negative, and negative otherwise. To assess significance, we grouped genes according to X variable, and then within each group we generated a bootstrap dataset. We estimated *ω*_*a*_, *ω*_*na*_, the mean value of X and Z for each group and the observed and predicted correlations between *ω*_*a*_, *ω*_*na*_, mean X and mean Z. We tabulated the number of bootstrap replicates in which predicted *r*_*YX*_ /observed *r*_*YX*_ > 1. We performed 100 bootstrap replicates for each analysis.

### Gene function analysis

Genes were divided by GO category and rates of adaptive and non-adaptive evolution were estimated for each category (note genes can contribute to multiple categories). For the VIP analysis we split each GO category into two groups – VIP and non-VIP genes, as per (Enard et al. 2016). To test whether there was significant variation in *ω*_*a*_ and *ω*_*na*_ across GO categories we shuffled data between gene labels; i.e. for each gene we have its synonymous and non-synonymous site frequency spectra and numbers of synonymous and non-synonymous substitutions. This data was randomly assigned to gene labels, hence preserving the covariance structure of the data -i.e. the fact that a gene can contribute to multiple GO categories. This shuffling was performed 100 times, each time recalculating *ω*_*a*_ and *ω*_*na*_.

We are interested in the extent to which the rate of adaptive and non-adaptive evolution is determined by whether it is a VIP gene versus other GO categorisations. We can quantify this by partitioning the variance in a two-way analysis of variance where the dimensions are VIP/non-VIP, and GO category. However, to estimate the variances we need to balance the data so that the error variance is the same for all cells in the two-way ANOVA. We did this by downsampling the data using a hypergeometric distribution, such that each cell had 200,000 combined non-synonymous and synonymous sites. To estimate the error variance we split the SFS and substitution data into two halves using a hypergeometric distribution and estimated *ω*_*a*_ and *ω*_*na*_ for each set; hence we have for each combination of VIP/non-VIP and GO category two estimates of the rate of adaptive and non-adaptive evolution, where the error variances for these estimates should be approximately equal.

## References

Albà, M. M., & Castresana, J. (2005). Inverse relationship between evolutionary rate and age of mammalian genes. Molecular Biology and Evolution, 22(3), 598–606. https://doi.org/10.1093/molbev/msi045

Arguello, J. R., Zhang, Y., Kado, T., Fan, C., Zhao, R., Innan, H., Wang, W., & Long, M. (2010). Recombination yet inefficient selection along the Drosophila melanogaster subgroup’s fourth chromosome. Molecular Biology and Evolution, 27(4), 848–861. https://doi.org/10.1093/molbev/msp291

Ashburner, M., Ball, C. A., Blake, J. A., Botstein, D., Butler, H., Cherry, J. M., Davis, A. P., Dolinski, K., Dwight, S. S., Eppig, J. T., Harris, M. A., Hill, D. P., Issel-Tarver, L., Kasarskis, A., Lewis, S., Matese, J. C., Richardson, J. E., Ringwald, M., Rubin, G. M., & Sherlock, G. (2000). Gene ontology: Tool for the unification of biology. The Gene Ontology Consortium. Nature Genetics, 25(1), 25–29. https://doi.org/10.1038/75556

Berglund, J., Pollard, K. S., & Webster, M. T. (2009). Hotspots of Biased Nucleotide Substitutions in Human Genes. PLoS Biology, 7(1), e1000026. https://doi.org/10.1371/journal.pbio.1000026

Betancourt, A. J., Welch, J. J., & Charlesworth, B. (2009). Reduced effectiveness of selection caused by a lack of recombination. Current Biology: CB, 19(8), 655–660. https://doi.org/10.1016/j.cub.2009.02.039

Bierne, N., & Eyre-Walker, A. (2004). The genomic rate of adaptive amino acid substitution in Drosophila. Molecular Biology and Evolution, 21(7), 1350–1360. https://doi.org/10.1093/molbev/msh134

Boyko, A. R., Williamson, S. H., Indap, A. R., Degenhardt, J. D., Hernandez, R. D., Lohmueller, K. E., Adams, M. D., Schmidt, S., Sninsky, J. J., Sunyaev, S. R., White, T. J., Nielsen, R., Clark, A. G., & Bustamante, C. D. (2008). Assessing the Evolutionary Impact of Amino Acid Mutations in the Human Genome. PLoS Genetics, 4(5), e1000083. https://doi.org/10.1371/journal.pgen.1000083

Burgess, R., & Yang, Z. (2008). Estimation of hominoid ancestral population sizes under bayesian coalescent models incorporating mutation rate variation and sequencing errors. Molecular Biology and Evolution, 25(9), 1979–1994. https://doi.org/10.1093/molbev/msn148

Bustamante, C. D., Fledel-Alon, A., Williamson, S., Nielsen, R., Todd Hubisz, M., Glanowski, S., Tanenbaum, D. M., White, T. J., Sninsky, J. J., Hernandez, R. D., Civello, D., Adams, M. D., Cargill, M., & Clark, A. G. (2005). Natural selection on protein-coding genes in the human genome. Nature, 437(7062), 1153–1157. https://doi.org/10.1038/nature04240

Cai, J. J., & Petrov, D. A. (2010). Relaxed Purifying Selection and Possibly High Rate of Adaptation in Primate Lineage-Specific Genes. Genome Biology and Evolution, 2, 393–409. https://doi.org/10.1093/gbe/evq019

Cai, J. J., Woo, P. C. Y., Lau, S. K. P., Smith, D. K., & Yuen, K.-Y. (2006). Accelerated evolutionary rate may be responsible for the emergence of lineage-specific genes in ascomycota. Journal of Molecular Evolution, 63(1), 1–11. https://doi.org/10.1007/s00239-004-0372-5

Campos, J. L., Halligan, D. L., Haddrill, P. R., & Charlesworth, B. (2014). The Relation between Recombination Rate and Patterns of Molecular Evolution and Variation in Drosophila melanogaster. Molecular Biology and Evolution, 31(4), 1010–1028. https://doi.org/10.1093/molbev/msu056

Castellano, D., Coronado-Zamora, M., Campos, J. L., Barbadilla, A., & Eyre-Walker, A. (2016). Adaptive Evolution Is Substantially Impeded by Hill-Robertson Interference in Drosophila. Molecular Biology and Evolution, 33(2), 442–455. https://doi.org/10.1093/molbev/msv236

Clark, A. G. (2003). Inferring Nonneutral Evolution from Human-Chimp-Mouse Orthologous Gene Trios. Science, 302(5652), 1960–1963. https://doi.org/10.1126/science.1088821

Cui, X., Lv, Y., Chen, M., Nikoloski, Z., Twell, D., & Zhang, D. (2015). Young Genes out of the Male: An Insight from Evolutionary Age Analysis of the Pollen Transcriptome. Molecular Plant, 8(6), 935–945. https://doi.org/10.1016/j.molp.2014.12.008

Daubin, V., & Ochman, H. (2004). Bacterial genomes as new gene homes: The genealogy of ORFans in E. coli. Genome Research, 14(6), 1036–1042. https://doi.org/10.1101/gr.2231904

Denamur, E., & Matic, I. (2006). Evolution of mutation rates in bacteria. Molecular Microbiology, 60(4), 820–827. https://doi.org/10.1111/j.1365-2958.2006.05150.x

Domazet-Loso, T., Brajkovic, J., & Tautz, D. (2007). A phylostratigraphy approach to uncover the genomic history of major adaptations in metazoan lineages. Trends in Genetics: TIG, 23(11), 533–539. https://doi.org/10.1016/j.tig.2007.08.014

Duret, L., & Galtier, N. (2009). Biased gene conversion and the evolution of mammalian genomic landscapes. Annual Review of Genomics and Human Genetics, 10, 285–311. https://doi.org/10.1146/annurev-genom-082908-150001

Edgar, R. C. (2004). MUSCLE: Multiple sequence alignment with high accuracy and high throughput. Nucleic Acids Research, 32(5), 1792–1797. https://doi.org/10.1093/nar/gkh340

Enard, D., Cai, L., Gwennap, C., & Petrov, D. A. (2016). Viruses are a dominant driver of protein adaptation in mammals. ELife, 5, e12469. https://doi.org/10.7554/eLife.12469

Eyre-Walker, A., & Keightley, P. D. (2009). Estimating the Rate of Adaptive Molecular Evolution in the Presence of Slightly Deleterious Mutations and Population Size Change. Molecular Biology and Evolution, 26(9), 2097–2108. https://doi.org/10.1093/molbev/msp119

Eyre-Walker, Adam. (2002). Changing effective population size and the McDonald-Kreitman test. Genetics, 162(4), 2017–2024.

Galtier, N. (2016). Adaptive Protein Evolution in Animals and the Effective Population Size Hypothesis. PLOS Genetics, 12(1), e1005774. https://doi.org/10.1371/journal.pgen.1005774

Galtier, N., & Duret, L. (2007). Adaptation or biased gene conversion? Extending the null hypothesis of molecular evolution. Trends in Genetics: TIG, 23(6), 273–277. https://doi.org/10.1016/j.tig.2007.03.011

Giraud, A. (2001). Costs and Benefits of High Mutation Rates: Adaptive Evolution of Bacteria in the Mouse Gut. Science, 291(5513), 2606–2608. https://doi.org/10.1126/science.1056421

Glémin, S. (2010). Surprising Fitness Consequences of GC-Biased Gene Conversion: I. Mutation Load and Inbreeding Depression. Genetics, 185(3), 939–959. https://doi.org/10.1534/genetics.110.116368

Gravel, S., Henn, B. M., Gutenkunst, R. N., Indap, A. R., Marth, G. T., Clark, A. G., Yu, F., Gibbs, R. A., The 1000 Genomes Project, Bustamante, C. D., Altshuler, D. L., Durbin, R. M., Abecasis, G. R., Bentley, D. R., Chakravarti, A., Clark, A. G., Collins, F. S., De La Vega, F. M., Donnelly, P., … McVean, G. A. (2011). Demographic history and rare allele sharing among human populations. Proceedings of the National Academy of Sciences, 108(29), 11983–11988. https://doi.org/10.1073/pnas.1019276108

The GTEx Consortium, Ardlie, K. G., Deluca, D. S., Segre, A. V., Sullivan, T. J., Young, T. R., Gelfand, E. T., Trowbridge, C. A., Maller, J. B., Tukiainen, T., Lek, M., Ward, L. D., Kheradpour, P., Iriarte, B., Meng, Y., Palmer, C. D., Esko, T., Winckler, W., Hirschhorn, J. N., … Dermitzakis, E. T. (2015). The Genotype-Tissue Expression (GTEx) pilot analysis: Multitissue gene regulation in humans. Science, 348(6235), 648–660. https://doi.org/10.1126/science.1262110

Haag-Liautard, C., Dorris, M., Maside, X., Macaskill, S., Halligan, D. L., Houle, D., Charlesworth, B., & Keightley, P. D. (2007). Direct estimation of per nucleotide and genomic deleterious mutation rates in Drosophila. Nature, 445(7123), 82–85. https://doi.org/10.1038/nature05388

Haerty, W., Jagadeeshan, S., Kulathinal, R. J., Wong, A., Ravi Ram, K., Sirot, L. K., Levesque, L., Artieri, C. G., Wolfner, M. F., Civetta, A., & Singh, R. S. (2007). Evolution in the fast lane: Rapidly evolving sex-related genes in Drosophila. Genetics, 177(3), 1321–1335. https://doi.org/10.1534/genetics.107.078865

Hill, W. G., & Robertson, A. (1966). The effect of linkage on limits to artificial selection. Genetical Research, 8(3), 269–294.

Hobolth, A., Christensen, O. F., Mailund, T., & Schierup, M. H. (2007). Genomic Relationships and Speciation Times of Human, Chimpanzee, and Gorilla Inferred from a Coalescent Hidden Markov Model. PLoS Genetics, 3(2), e7. https://doi.org/10.1371/journal.pgen.0030007

Hurst, L. D., & Smith, N. G. (1999). Do essential genes evolve slowly? Current Biology: CB, 9(14), 747–750. https://doi.org/10.1016/s0960-9822(99)80334-0

Krylov, D. M., Wolf, Y. I., Rogozin, I. B., & Koonin, E. V. (2003). Gene loss, protein sequence divergence, gene dispensability, expression level, and interactivity are correlated in eukaryotic evolution. Genome Research, 13(10), 2229–2235. https://doi.org/10.1101/gr.1589103

Lachance, J., & Tishkoff, S. A. (2014). Biased gene conversion skews allele frequencies in human populations, increasing the disease burden of recessive alleles. American Journal of Human Genetics, 95(4), 408–420. https://doi.org/10.1016/j.ajhg.2014.09.008

Lesecque, Y., Keightley, P. D., & Eyre-Walker, A. (2012). A resolution of the mutation load paradox in humans. Genetics, 191(4), 1321–1330. https://doi.org/10.1534/genetics.112.140343

Liao, B.-Y., Scott, N. M., & Zhang, J. (2006). Impacts of gene essentiality, expression pattern, and gene compactness on the evolutionary rate of mammalian proteins. Molecular Biology and Evolution, 23(11), 2072–2080. https://doi.org/10.1093/molbev/msl076

Lipman, D. J., Souvorov, A., Koonin, E. V., Panchenko, A. R., & Tatusova, T. A. (2002). The relationship of protein conservation and sequence length. BMC Evolutionary Biology, 2, 20. https://doi.org/10.1186/1471-2148-2-20

Litman, T., & Stein, W. D. (2019). Obtaining estimates for the ages of all the protein-coding genes and most of the ontology-identified noncoding genes of the human genome, assigned to 19 phylostrata. Seminars in Oncology, 46(1), 3–9. https://doi.org/10.1053/j.seminoncol.2018.11.002

Lohmueller, K. E., Indap, A. R., Schmidt, S., Boyko, A. R., Hernandez, R. D., Hubisz, M. J., Sninsky, J. J., White, T. J., Sunyaev, S. R., Nielsen, R., Clark, A. G., & Bustamante, C. D. (2008). Proportionally more deleterious genetic variation in European than in African populations. Nature, 451(7181), 994–997. https://doi.org/10.1038/nature06611

Long, M., & Langley, C. (1993). Natural selection and the origin of jingwei, a chimeric processed functional gene in Drosophila. Science, 260(5104), 91–95. https://doi.org/10.1126/science.7682012

Lynch, M. (2002). GENOMICS: Gene Duplication and Evolution. Science, 297(5583), 945–947. https://doi.org/10.1126/science.1075472

Lynch, Michael, Ackerman, M. S., Gout, J.-F., Long, H., Sung, W., Thomas, W. K., & Foster, P. L. (2016). Genetic drift, selection and the evolution of the mutation rate. Nature Reviews Genetics, 17(11), 704–714. https://doi.org/10.1038/nrg.2016.104

Mackay, T. F. C., Richards, S., Stone, E. A., Barbadilla, A., Ayroles, J. F., Zhu, D., Casillas, S., Han, Y., Magwire, M. M., Cridland, J. M., Richardson, M. F., Anholt, R. R. H., Barrón, M., Bess, C., Blankenburg, K. P., Carbone, M. A., Castellano, D., Chaboub, L., Duncan, L., … Gibbs, R. A. (2012). The Drosophila melanogaster Genetic Reference Panel. Nature, 482(7384), 173–178. https://doi.org/10.1038/nature10811

Marais, G., & Charlesworth, B. (2003). Genome evolution: Recombination speeds up adaptive evolution. Current Biology: CB, 13(2), R68–70. https://doi.org/10.1016/s0960-9822(02)01432-x

McDonald, J. H., & Kreitman, M. (1991). Adaptive protein evolution at the Adh locus in Drosophila. Nature, 351(6328), 652–654. https://doi.org/10.1038/351652a0

Messer, P. W., & Petrov, D. A. (2013). Frequent adaptation and the McDonald-Kreitman test. Proceedings of the National Academy of Sciences of the United States of America, 110(21), 8615–8620. https://doi.org/10.1073/pnas.1220835110

Moutinho, A. F., Trancoso, F. F., & Dutheil, J. Y. (2019). The Impact of Protein Architecture on Adaptive Evolution. Molecular Biology and Evolution, 36(9), 2013–2028. https://doi.org/10.1093/molbev/msz134

Necşulea, A., Popa, A., Cooper, D. N., Stenson, P. D., Mouchiroud, D., Gautier, C., & Duret, L. (2011). Meiotic recombination favors the spreading of deleterious mutations in human populations. Human Mutation, 32(2), 198–206. https://doi.org/10.1002/humu.21407

Neme, R., & Tautz, D. (2013). Phylogenetic patterns of emergence of new genes support a model of frequent de novo evolution. BMC Genomics, 14, 117. https://doi.org/10.1186/1471-2164-14-117

Nielsen, R., Bustamante, C., Clark, A. G., Glanowski, S., Sackton, T. B., Hubisz, M. J., Fledel-Alon, A., Tanenbaum, D. M., Civello, D., White, T. J., J. Sninsky J,. Adams, M. D., & Cargill, M. (2005). A Scan for Positively Selected Genes in the Genomes of Humans and Chimpanzees. PLoS Biology, 3(6), e170. https://doi.org/10.1371/journal.pbio.0030170

Obbard, D. J., Welch, J. J., Kim, K.-W., & Jiggins, F. M. (2009). Quantifying Adaptive Evolution in the Drosophila Immune System. PLoS Genetics, 5(10), e1000698. https://doi.org/10.1371/journal.pgen.1000698

Pál, C., Papp, B., & Hurst, L. D. (2001). Highly expressed genes in yeast evolve slowly. Genetics, 158(2), 927–931.

Papatheodorou, I., Moreno, P., Manning, J., Fuentes, A. M.-P., George, N., Fexova, S., Fonseca, N. A., Füllgrabe, A., Green, M., Huang, N., Huerta, L., Iqbal, H., Jianu, M., Mohammed, S., Zhao, L., Jarnuczak, A. F., Jupp, S., Marioni, J., Meyer, K., … Brazma, A. (2019). Expression Atlas update: From tissues to single cells. Nucleic Acids Research, gkz947. https://doi.org/10.1093/nar/gkz947

Prado-Martinez, J., Sudmant, P. H., Kidd, J. M., Li, H., Kelley, J. L., Lorente-Galdos, B., Veeramah, K. R., Woerner, A. E., O’Connor, T. D., Santpere, G., Cagan, A., Theunert, C., Casals, F., Laayouni, H., Munch, K., Hobolth, A., Halager, A. E., Malig, M., Hernandez-Rodriguez, J., … Marques-Bonet, T. (2013). Great ape genetic diversity and population history. Nature, 499(7459), 471–475. https://doi.org/10.1038/nature12228

Presgraves, D. C. (2005). Recombination enhances protein adaptation in Drosophila melanogaster. Current Biology: CB, 15(18), 1651–1656. https://doi.org/10.1016/j.cub.2005.07.065

Pröschel, M., Zhang, Z., & Parsch, J. (2006). Widespread adaptive evolution of Drosophila genes with sex-biased expression. Genetics, 174(2), 893–900. https://doi.org/10.1534/genetics.106.058008

Ratnakumar, A., Mousset, S., Glémin, S., Berglund, J., Galtier, N., Duret, L., & Webster, M. T. (2010). Detecting positive selection within genomes: The problem of biased gene conversion. Philosophical Transactions of the Royal Society of London. Series B, Biological Sciences, 365(1552), 2571–2580. https://doi.org/10.1098/rstb.2010.0007

Rocha, E. P. C., & Danchin, A. (2004). An analysis of determinants of amino acids substitution rates in bacterial proteins. Molecular Biology and Evolution, 21(1), 108–116. https://doi.org/10.1093/molbev/msh004

Rousselle, M., Simion, P., Tilak, M.-K., Figuet, E., Nabholz, B., & Galtier, N. (2020). Is adaptation limited by mutation? A timescale-dependent effect of genetic diversity on the adaptive substitution rate in animals. PLOS Genetics, 16(4), e1008668. https://doi.org/10.1371/journal.pgen.1008668

Sackton, T. B., Lazzaro, B. P., Schlenke, T. A., Evans, J. D., Hultmark, D., & Clark, A. G. (2007). Dynamic evolution of the innate immune system in Drosophila. Nature Genetics, 39(12), 1461–1468. https://doi.org/10.1038/ng.2007.60

Schrago, C. G. (2014). The Effective Population Sizes of the Anthropoid Ancestors of the Human-Chimpanzee Lineage Provide Insights on the Historical Biogeography of the Great Apes. Molecular Biology and Evolution, 31(1), 37–47. https://doi.org/10.1093/molbev/mst191

Soni, V., Eyre-Walker, A. (2021). Site level factors that affect the rate of adaptive evolution in humans and chimpanzees; the effect of contracting population size. (Unpublished).

Spence, J. P., & Song, Y. S. (2019). Inference and analysis of population-specific fine-scale recombination maps across 26 diverse human populations. Science Advances, 5(10), eaaw9206. https://doi.org/10.1126/sciadv.aaw9206

Subramanian, S., & Kumar, S. (2004). Gene expression intensity shapes evolutionary rates of the proteins encoded by the vertebrate genome. Genetics, 168(1), 373–381. https://doi.org/10.1534/genetics.104.028944

Taddei, F., Radman, M., Maynard-Smith, J., Toupance, B., Gouyon, P. H., & Godelle, B. (1997). Role of mutator alleles in adaptive evolution. Nature, 387(6634), 700–702. https://doi.org/10.1038/42696

Tautz, D., & Domazet-Lošo, T. (2011). The evolutionary origin of orphan genes. Nature Reviews. Genetics, 12(10), 692–702. https://doi.org/10.1038/nrg3053

Tenaillon, O., Toupance, B., Le Nagard, H., Taddei, F., & Godelle, B. (1999). Mutators, population size, adaptive landscape and the adaptation of asexual populations of bacteria. Genetics, 152(2), 485–493.

The 1000 Genomes Project Consortium. (2015). A global reference for human genetic variation. Nature, 526(7571), 68–74. https://doi.org/10.1038/nature15393

The Chimpanzee Sequencing and Analysis Consortium. (2005). Initial sequence of the chimpanzee genome and comparison with the human genome. Nature, 437(7055), 69–87. https://doi.org/10.1038/nature04072

The Gene Ontology Consortium, Carbon, S., Douglass, E., Good, B. M., Unni, D. R., Harris, N. L., Mungall, C. J., Basu, S., Chisholm, R. L., Dodson, R. J., Hartline, E., Fey, P., Thomas, P. D., Albou, L.-P., Ebert, D., Kesling, M. J., Mi, H., Muruganujan, A., Huang, X., … Elser, J. (2021). The Gene Ontology resource: Enriching a GOld mine. Nucleic Acids Research, 49(D1), D325–D334. https://doi.org/10.1093/nar/gkaa1113

Thornton, K., & Long, M. (2002). Rapid divergence of gene duplicates on the Drosophila melanogaster X chromosome. Molecular Biology and Evolution, 19(6), 918–925. https://doi.org/10.1093/oxfordjournals.molbev.a004149

Vishnoi, A., Kryazhimskiy, S., Bazykin, G. A., Hannenhalli, S., & Plotkin, J. B. (2010). Young proteins experience more variable selection pressures than old proteins. Genome Research, 20(11), 1574–1581. https://doi.org/10.1101/gr.109595.110

Wang, W., Zheng, H., Yang, S., Yu, H., Li, J., Jiang, H., Su, J., Yang, L., Zhang, J., McDermott, J., Samudrala, R., Wang, J., Yang, H., Yu, J., Kristiansen, K., Wong, G. K.-S., & Wang, J. (2005). Origin and evolution of new exons in rodents. Genome Research, 15(9), 1258–1264. https://doi.org/10.1101/gr.3929705

Wolf, Y. I., Novichkov, P. S., Karev, G. P., Koonin, E. V., & Lipman, D. J. (2009). The universal distribution of evolutionary rates of genes and distinct characteristics of eukaryotic genes of different apparent ages. Proceedings of the National Academy of Sciences of the United States of America, 106(18), 7273–7280. https://doi.org/10.1073/pnas.0901808106

Wright, S. (1931). Evolution in Mendelian Populations. Genetics, 16(2), 97–159.

Wright, S. (1932). The roles of mutation, inbreeding, crossbreeding and selection in evolution. Sixth Int. Congr. Genet. 1:356–366

Yang, Z. (2007). PAML 4: Phylogenetic Analysis by Maximum Likelihood. Molecular Biology and Evolution, 24(8), 1586–1591. https://doi.org/10.1093/molbev/msm088

Yates, A. D., Achuthan, P., Akanni, W., Allen, J., Allen, J., Alvarez-Jarreta, J., Amode, M. R., Armean, I. M., Azov, A. G., Bennett, R., Bhai, J., Billis, K., Boddu, S., Marugán, J. C., Cummins, C., Davidson, C., Dodiya, K., Fatima, R., Gall, A., … Flicek, P. (2019). Ensembl 2020. Nucleic Acids Research, gkz966. https://doi.org/10.1093/nar/gkz966

Zhang, J. (2000). Protein-length distributions for the three domains of life. Trends in Genetics: TIG, 16(3), 107–109. https://doi.org/10.1016/s0168-9525(99)01922-8

Zhang, Y. E., Vibranovski, M. D., Krinsky, B. H., & Long, M. (2010). Age-dependent chromosomal distribution of male-biased genes in Drosophila. Genome Research, 20(11), 1526–1533. https://doi.org/10.1101/gr.107334.110

