## supplementary material for "Factors that affect the rates of adaptive and non-adaptive evolution at the gene level in humans and chimpanzees"

| | | | Both fixed | Both random |
|---|---|---|---|---|
| **Term** | **Number** | **MS** | **Variance** | **Variance** |
| Residual | 2 | 5.2E-05 | 0.000052 | 0.000052 |
| VIP | 2 | 0.02919 | 0.00112054 | 0.001120538 |
| GO | 13 | 0.00097 | 0.000229 | 0.000229 |

**Supplementary table S1:** Estimated variance components from two-way analysis of variance on $\omega_a$ for GO categories with 200,000 sites or more.

| Term | Number | MS | Both fixed Variance | Both random Variance |
|---|---|---|---|---|
| Residual | 2 | 0.000017 | 0.000017 | 0.000017 |
| VIP | 2 | 0.000983 | 3.7154E-05 | 3.71538E-05 |
| GO | 13 | 0.000034 | 4.25E-6 | 4.25E-06 |

**Supplementary table S2:** Estimated variance components from two-way analysis of variance on $\omega_{na}$ for GO categories with 200,000 sites or more.

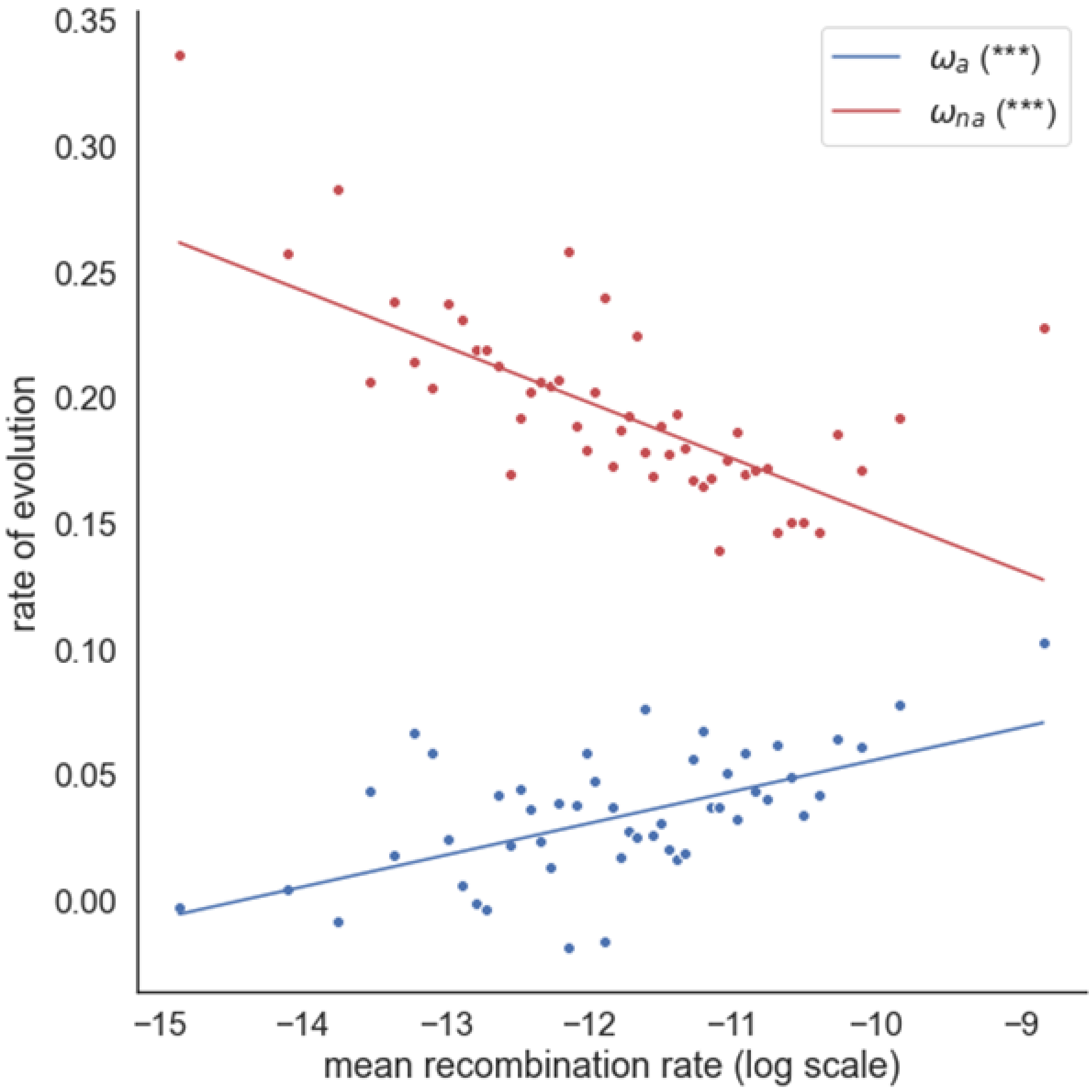


**Supplementary figure S1:** Estimates of $\omega_a$ and $\omega_{na}$ plotted against the log of the mean recombination rate for genes binned into 50 recombination bins of equal size. An unweighted linear regression is fitted to the data. The significance of each correlation is shown in the plot legend, (*P < 0.05; **P < 0.01; ***P < 0.001; "." 0.05 ≤ P < 0.10).

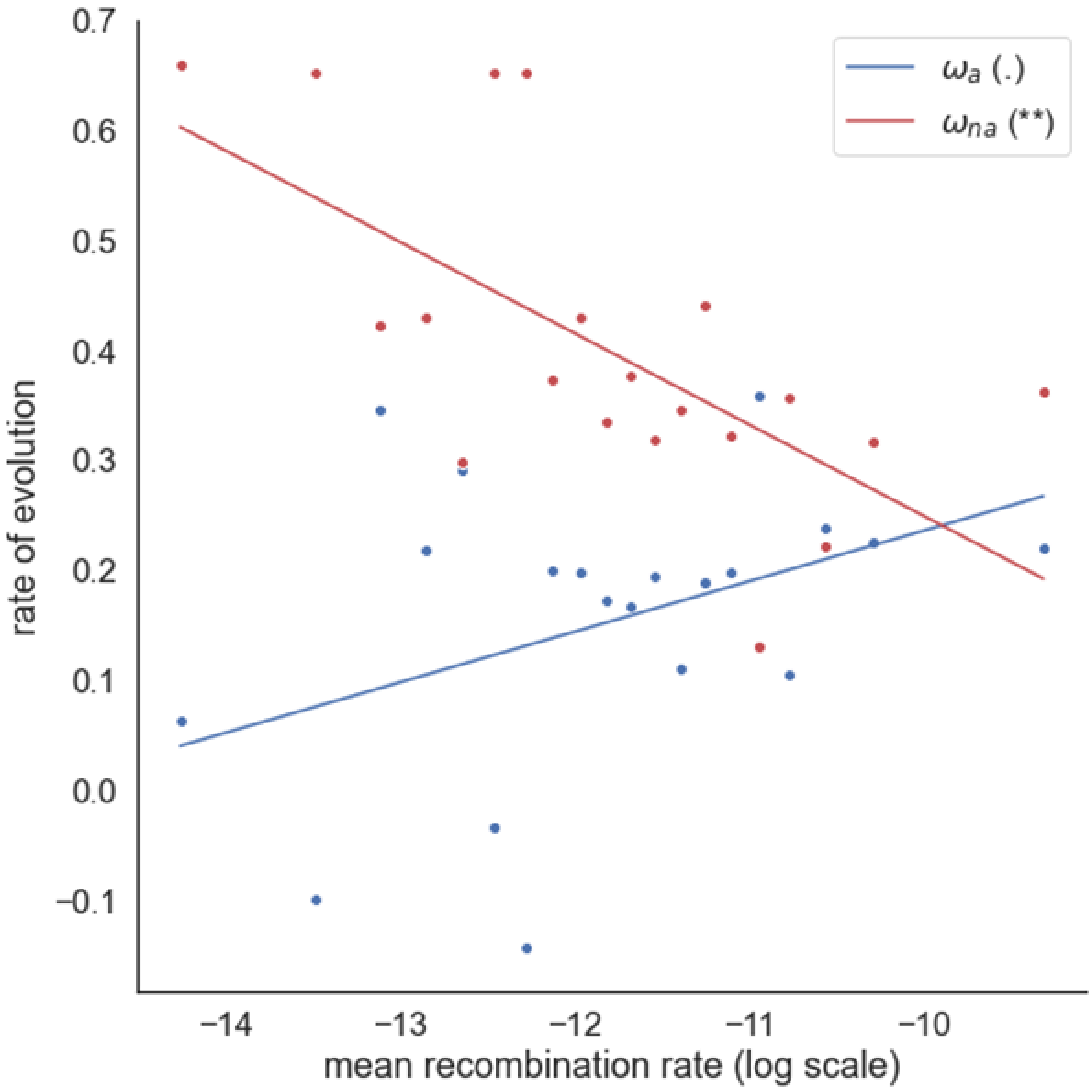


**Supplementary figure S2:** Estimates of $\omega_a$ and $\omega_{na}$ plotted against the log of the mean recombination rate, controlling for biased gene conversion, for genes binned into 20 recombination bins of equal size. An unweighted linear regression is fitted to the data. The significance of each correlation is shown in the plot legend, (*P < 0.05; **P < 0.01; ***P < 0.001; "." 0.05 ≤ P < 0.10).

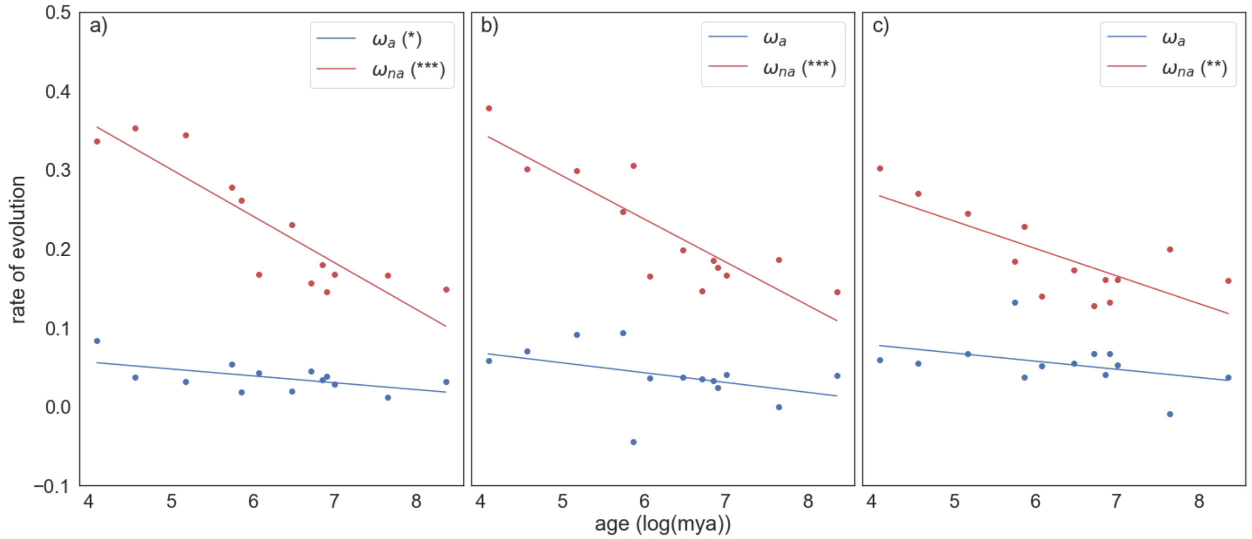


**Supplementary figure S3:** Estimates of $\omega_a$ and $\omega_{na}$ plotted against log gene age for genes binned into phylostratigraphic age categories, controlling for a) recombination rate; b) gene length; c) gene expression. An unweighted linear regression is fitted to the data. The significance of each correlation is shown in the plot legend, (*$P < 0.05$; **$P < 0.01$; ***$P < 0.001$; "." $0.05 \leq P < 0.10$).

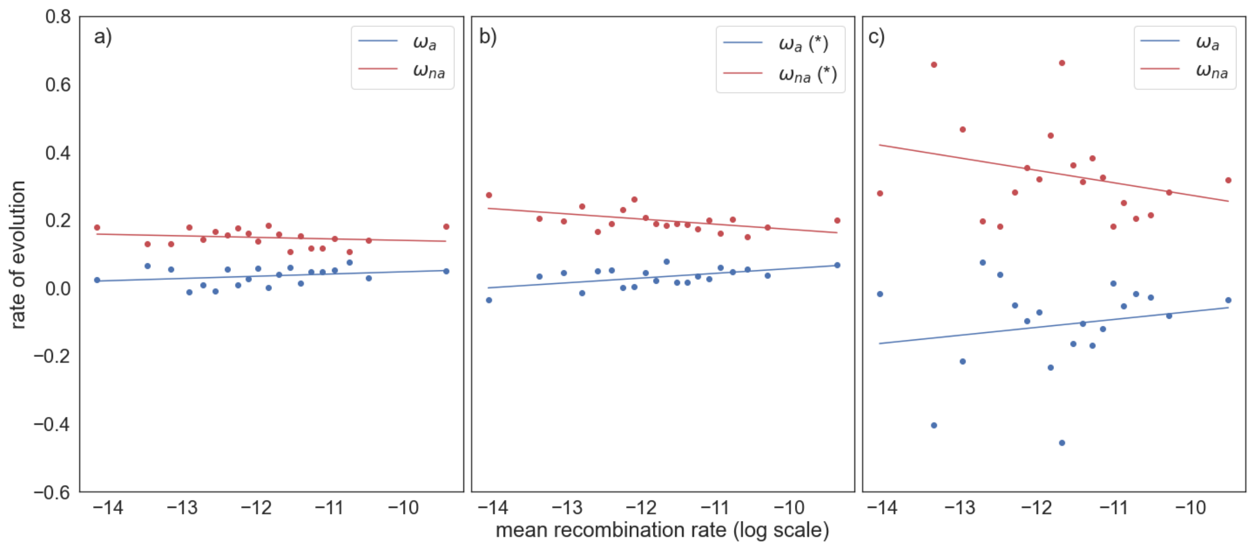


**Supplementary figure S4:** Estimates of $\omega_a$ and $\omega_{na}$ plotted against the log of the mean recombination rate for genes binned into 20 recombination bins of equal size, controlling for a) gene age; b) gene length; c) gene expression. An unweighted linear regression is fitted to the data. The significance of each correlation is shown in the plot legend, (*P < 0.05; **P < 0.01; ***P < 0.001; "." 0.05 ≤ P < 0.10).

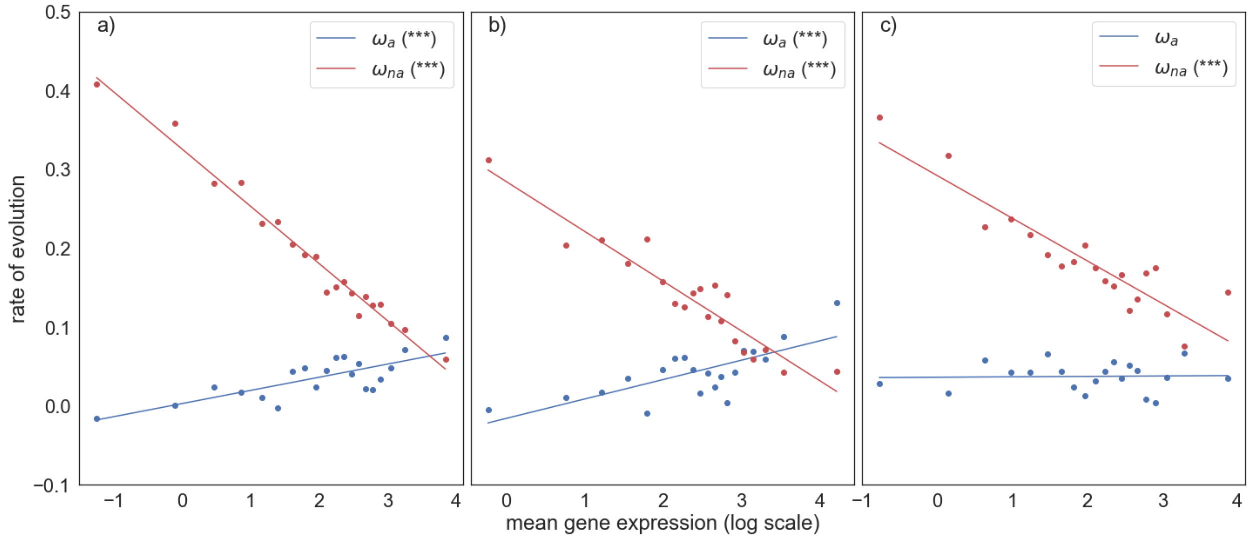


**Supplementary figure S5:** Estimates of $\omega_a$ and $\omega_{na}$ plotted against the log of the mean gene expression for genes binned into 20 mean expression bins of equal size, controlling for a) recombination rate; b) gene age; c) gene length. An unweighted linear regression is fitted to the data. The significance of each correlation is shown in the plot legend, (*P < 0.05; **P < 0.01; ***P < 0.001; "." 0.05 ≤ P < 0.10).

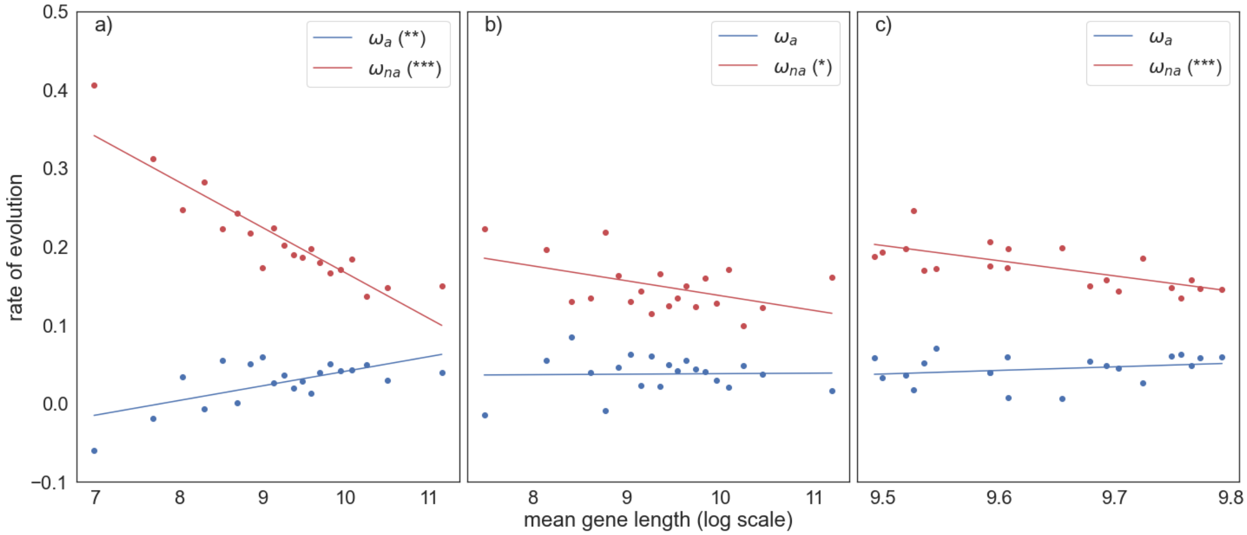


**Supplementary figure S6:** Estimates of $\omega_a$ and $\omega_{na}$ plotted against the log of the mean gene length for genes binned into 20 mean length bins of equal size, controlling for a) recombination rate; b) gene age; c) gene expression. An unweighted linear regression is fitted to the data. The significance of each correlation is shown in the plot legend, (*P < 0.05; **P < 0.01; ***P < 0.001; "." 0.05 ≤ P < 0.10).
